# Eco-evolutionary dynamics in two-species mutualistic systems: One-sided population decline triggers joint interaction disinvestment

**DOI:** 10.1101/2023.02.12.528164

**Authors:** Franz Weyerer, Avril Weinbach, Christiane Zarfl, Korinna T. Allhoff

**Affiliations:** Eberhard Karls Universität Tübingen, Tübingen, Germany; Universität Hohenheim, Stuttgart, Germany

**Keywords:** Evolutionary rescue, Evolutionary murder, Coevolution, Mutualism, Insect decline, Adaptive Dynamics

## Abstract

The interplay between ecological and evolutionary dynamics can create feedback that reinforces external disturbances and potentially threatens species’ coexistence. For example, plants might invest less into attracting insect pollinators (decreased flower or nectar production) and more into independence (increased selfing or vegetative reproduction) when faced with pollinator decline. This adaptive response saves plant resources at the cost of further threatening the pollinator population. We ask under which conditions such self-reinforcing feedback occurs in two-species mutualistic systems when considering one-sided population decline and whether it can be counteracted by self-dampening feedback if co-evolution of both interaction partners is considered. Based on a mathematical model and in line with previous studies, we find that the described pattern of accelerated population decline occurs for a wide range of parameter values if a concave allocation trade-off between independent growth and interaction investment is assumed. The undisturbed population typically disinvests first, which then forces the declining population to also disinvest, in favour of other energy sources. However, a decelerated population decline can occur if the adaptation of the undisturbed partner is relatively slow compared to environmental decay, reducing the speed of its disinvestment, or if the initial investment into the interaction was very high. Our results suggest that if actions are taken to save endangered populations, not only the evolution of the target species but also of their interaction partner, as well as the interaction between them should be considered.

## 1. Introduction

Insects experience stressors from many directions (Wagner et al., 2021), be it the fragmentation, degradation, and destruction of habitats (Fabienne Harris & Johnson, 2004), the enhanced use of pesticides (Hoffmann et al., 2010), invasive species (Madjidian et al., 2008), new diseases (Le Conte et al., 2010) or the effects of climate change in general. All these factors have led to a decline in the diversity and biomass of flying insects (Fox, 2013; Hallmann et al., 2017; Potts et al., 2010). This is not only a problem for the insect populations themselves but also for all species that interact with them, either antagonistically or mutualistically. Mutualistic interactions, which play a crucial role in sustaining ecosystem functioning and services (Reid et al., 2019), are of special importance in this context, given that they are less stable than antagonistic ones and more likely to be lost (Sachs & Simms, 2006; Toby Kiers et al., 2010).

The stressors mentioned above put insect populations under pressure to adjust to new environmental conditions, e.g. via phenotypic plasticity or adaptation, which in turn forces their mutualistic interaction partners to adjust as well. In a worst-case scenario, one species may adapt too slowly to catch up with the interaction partner and their dependence on each other could then lead to co-extinction. In other cases, the cost of maintaining the mutualistic interaction might simply increase under the new environmental conditions, meaning that the interaction itself could become subject to change in case the costs ultimately outweigh the benefits (Bronstein, 2001). This can lead to a mutualism loss, a partner switch, or even change the type of interaction from mutualism to antagonism (Toby Kiers et al., 2010).

Adaptive and evolutionary responses to external stressors are not limited to timescales of hundreds or thousands of years but can occur surprisingly fast (Cheptou et al., 2008; Hairston et al., 2005; Thompson, 1998). For example, Roels and Kelly (2011) showed that plants (*Mimulus guttatus*) increased their self autonomous reproduction in response to loss of pollinators (*Bombus impatiens*) within only a few generations. This trend is typically associated with smaller flowers, reduced flower longevity, and fewer rewards for the pollinator (Thomann et al., 2013). On the other hand, Gervasi and Schiestl (2017) showed that after exposing plants (*Brassica rapa*) to pollinators (*Bombus terrestris)*, the plants evolved to higher investment in plant height and amount of volatile emission to attract the pollinator. Both plant responses potentially reinforce the initial change in pollinator abundance by decreasing or increasing food resources, creating so-called eco-evolutionary feedback (Pelletier et al., 2009; Post & Palkovacs, 2009).

Quantifying the impact of such eco-evolutionary feedback on system dynamics and species survival is very challenging, especially in systems that involve more than one species (Koch et al., 2014; N. Loeuille, 2019), but the mathematical framework of adaptive dynamics can help to gain valuable insights (Brännström et al., 2013; Dieckmann & Law, 1996; Ferriere & Legendre, 2013; Geritz et al., 1998; Govaert et al., 2019). For instance, Weinbach et al. (2022) used this method to investigate eco-evolutionary dynamics in a simple Lotka-Volterra mutualism system. They studied this system through the lens of plant-pollinator interactions. One of the interaction partners, which was interpreted as the pollinator, experienced a sudden decline in intrinsic growth due to external stressors. The other interaction partner, which was interpreted as a plant, was then able to react to changes in pollinator abundances by shifting its energy allocation. It could either invest into attractiveness for the pollinator, in order to maintain the interaction, or into its own intrinsic growth, in order to become more independent. The model suggests that plants tend to disinvest from the interaction in the face of pollinator decline, in line with the empirical evidence from Roels and Kelly (2011) mentioned above, if a concave trade-off function was assumed. This disinvestment could then further threaten the already declining pollinator population, resulting in self-reinforcing eco-evolutionary feedback. The authors hypothesised that plant disinvestment could even result in so-called eco-evolutionary murder (name suggested by Parvinen in 2005), meaning that the pollinator does not go extinct as a direct consequence of an external stressor but due to plant evolution.

It remains unclear, however, whether this key finding is robust to changes in the underlying model assumptions. This is in particular relevant when considering co-evolution of both interaction partners. We hypothesise that if pollinators were also allowed to re-adjust their energy investment in response to changing conditions, similar to the plants, then this might counteract the impact of external stressors. In consequence, we would observe self-dampening instead of self-reinforcing eco-evolutionary feedback and hence decelerated instead of accelerated pollinator decline. Pollinator evolution could even enable so-called evolutionary rescue (S. M. Carlson et al., 2014; Ferriere & Legendre, 2013, 2013; Gomulkiewicz & Holt, 1995; Gonzalez et al., 2013), meaning that pollinators are able to survive environmental conditions that would lead to extinction in the absence of evolution.

To test this hypothesis, we extended the model by Weinbach et al. to account for coevolution of both interaction partners. More specifically, we analysed the evolution of the investments in the mutualistic interaction of both species in response to the declining abundance of one interaction partner. By design, the declining interaction partner sooner or later reached the extinction threshold. We asked under which conditions the self-reinforcing feedback found by Weinbach et al. (2022) dominates system dynamics, resulting in earlier extinction, and what conditions have to be met to observe the reversed outcome, that is delayed extinction. To address these questions, we varied ecological model parameters, in particular interaction benefits and intraspecific competition, as well as the relative timescales of the environmental decay versus the evolutionary responses of both interaction partners.

## 2. Model & Methods

In the following, we first give an overview of the ecological processes considered in the model before explaining how population decline due to environmental decay is implemented. We then specify how evolutionary changes in energy investment are taken into account and how all three model components (ecology, environment and evolution) are linked within our numerical algorithm. A summary of all model parameters can be found in Table 1.

**Table 1:** Overview of model parameters. Base values correspond to the initial eco-evolutionary equilibrium used in reference simulation S1.

| Definition | Symbol | Unit | Base value | Analysed range |
| --- | --- | --- | --- | --- |
| **Ecological parameter** |  |  |  |  |
| Plant net growth coefficient | $r_P$ | 1/time | 0.66 | 0 – 1 |
| Pollinator net growth coefficient | $r_A$ | 1/time | 0.66 | 0 – 1 |
| Pollinator growth without env. decay | $r'_A$ | 1/time | 0.66 | <1 |
| Plant intraspecific competition | $c_P$ | m<sup>2</sup>/time | 1.00 | 0.9–2.5 |
| Pollinator intraspecific competition | $c_A$ | m<sup>2</sup>/time | 1.00 | 0.9–2.5 |
| Plant interaction investment | $\alpha$ | 1 | 0.72 | 0 – 0.80 |
| Pollinator interaction investment | $\beta$ | 1 | 0.72 | 0 – 0.80 |
| Plant interaction benefit provided by the pollinator | $\gamma_A$ | m<sup>2</sup>/time | 1.4 | 0 – 1.50 |
| Pollinator interaction benefit provided by the plant | $\gamma_P$ | m<sup>2</sup>/time | 1.4 | 0 – 1.50 |
| **Environmental parameter** |  |  |  |  |
| Environmental decay constant | $\delta_{Env}$ | 1/time<sup>2</sup> | 0.001 | 0.1-0.00001 |
| Extinction threshold | $\varepsilon$ | 1/m<sup>2</sup> | 0.01 | 0.01 |
| **Evolutionary parameter** |  |  |  |  |
| Plant mutation probability | $q_P$ | [%] | 0 | 0 – 100 |
| Pollinator mutation probability | $q_A$ | [%] | 0 | 0 – 100 |
| Plant mutation maximum step size | $\Delta\alpha$ | 1 | 0.016 | 0.016 |
| Pollinator mutation maximum step size | $\Delta\beta$ | 1 | 0.016 | 0.016 |
| Trade-off shape | $s$ | 1 | 3 | 3 |

### 2.1 Ecological dynamics

As in Weinbach et al. (2022) the basic model consists of two interacting species. The model is relatively abstract and not system-specific, meaning that it could in principle describe any mutualistic system (e.g. plant-mycorrhizal, plant-ant, or ant-fungi systems, see Bronstein (1994)). However, we refer to the two interaction partners as plant population *P* and pollinator population *A* [1/m^2^]. The population dynamics are given by a Lotka-Volterra-type model with two differential equations:

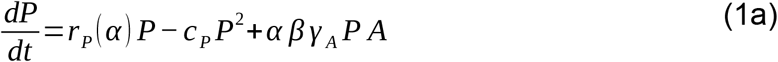

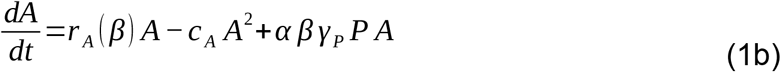

The equations consist of three terms each. The first term describes the independent growth of the populations, where *r*_*P*_ and *r*_*A*_ represent the net growth coefficients in [1/time] of the plant and the pollinator, respectively. The plant coefficient includes processes like selfing or vegetative reproduction that allow for growth even in the absence of pollinators. The pollinator coefficient, on the other hand, includes not only energetic gain from alternative food sources, reproduction spots or shelter, but also mortality losses. It can become negative, if those losses outweigh the positive terms. Both coefficients are dependent on the corresponding interaction parameters *α* or *β* via trade-off functions, as explained in more detail further below.

The second term describes the intraspecific competition for both plants and pollinators, with coefficients *c*_*P*_ and *c*_*A*_ in [m^2^/time], respectively. Competition is assumed to be at a constant level, meaning that evolution and also environmental decay do not change the intensity of the competition, in line with the original model by Weinbach et al. (2022). The first two terms combined give logistic growth with carrying capacities *r*_*i*_*/c*_*i*_ for the plant (*i=P*) and the pollinator *(i=A*).

The last term finally defines the mutualistic interaction between plants and pollinators. More precisely, *γ* _*A*_ (measured in [m^2^/time]) describes the interaction benefits for the plant (e.g. sexual reproduction due to the fertilisation or protection against herbivory provided by the mutualist interactor), while *γ* _*P*_ corresponds to the benefits for the pollinator (e.g. energetic gain due to nectar, pollen, or other plant exudates provided by the plant). The parameters *α* and *β* are dimensionless and capture the investment of the plant and the pollinator into the interaction. On the plant side, this might describe the amount of nectar or size, shape, colour or number of flowers (Willmer, 2011). On the pollinator side, this could represent the pollinator’s morphology, such as adapted mouthparts, or the designated time to search for this particular plant. Note that without *β* the model simplifies into the original model version by Weinbach et al. (2022). Also note that we chose a linear functional response mainly for the purpose of consistency with Weinbach et al. (2022). This choice has the advantage of keeping the model itself rather general and relatively simple, which allows for explicit analytical investigations.

The basic ecological dynamics of the model have already been investigated by Weinbach et al. (2022), but the results have to be adjusted to the introduction of *β*, as explained in the following. A stable equilibrium for ecological coexistence can be found at the crossing point of the non-trivial zero-growth isoclines (see online supplementary material A for more details). The corresponding equilibrium population densities can be calculated via the following equations:

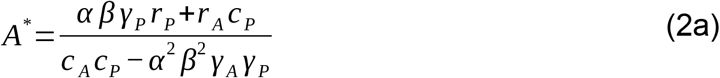

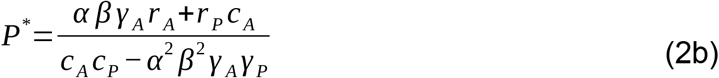

The equilibrium point is only feasible for positive population densities. This condition is fulfilled, assuming that all parameters are positive, if the denominator is positive:

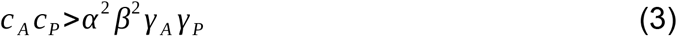

The above inequality can be interpreted as a balance between intraspecific competition (left) and interspecific mutualism (right). If the condition of inequality (3) is not fulfilled, the self-regulatory effect of the intraspecific competition is too weak to compensate for the positive feedback that arises from the mutualistic interaction, resulting in unbounded population growth. Such unlimited growth is usually controlled by additional processes, such as predation or interspecific competition, which are not explicitly considered in the model. Instead, Weinbach et al. defined a critical level for plant interaction investment *α*_*cl*_. The same is done here for the critical level of pollinator interaction investment *β*_*cl*_:

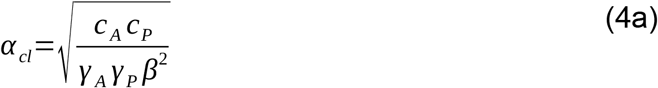

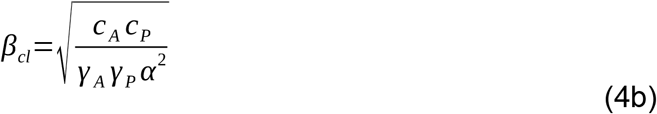

Stable coexistence of both populations is thus only possible if 0 *≤α ≤α* _*max*_<*α* _*cl*_ and 0 *≤ β ≤ β*_*max*_ < *β*_*cl*_. In consistency with Weinbach et al., we set *α*_*max*_=0.8 * *α*_*cl*_. To keep the existing symmetry between both populations, we furthermore set *β*_*max*_=0.8 * *β*_*cl*_.

### 2.2 Environmental decay

We assume a steady decline in habitat quality, which directly affects the mortality of pollinators, so that the net growth rates of pollinators continuously decrease. We furthermore assume that this decrease can be described via the following equation:

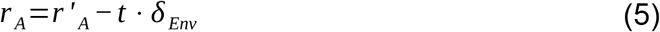

The parameters *r ‘*_*A*_ and *δ*_*Env*_ denote the initial growth rate at time t=0 and the change in pollinator mortality rate per unit time, respectively. Note that *r* _*A*_= *r ‘* _*A*_ in the absence of environmental decline, that is when *δ*_*Env*_= 0. In all other cases, we can expect that the habitat quality will eventually become so bad that pollinators are no longer able to survive. Simulations are stopped as soon as the pollinator population density falls below the extinction threshold *ε*.

### 2.3 Evolution of interaction investment

Both plants and pollinators are assumed to have a finite amount of energy that they can invest either into independence from each other (via the growth rates *r*_*P*_ or *r’*_*A*_) or into the joint mutualistic interaction (via the investment into the interaction *α* or *β*). We use the same trade-off function that Weinbach et al. (2022) introduced for the plant also for the pollinator:

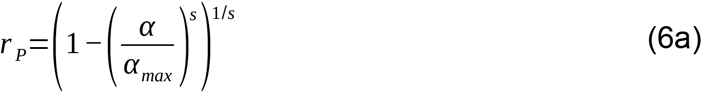

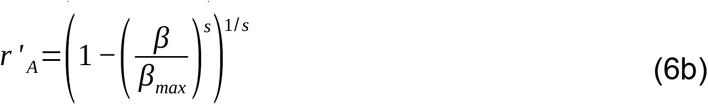

The exponent *s* in these trade-off functions controls the trade-off shape. Weinbach et al. (2022) found that the plant evolution strongly depends on this parameter. Small values of *s* translate into a convex trade-off shape (see supplementary Fig. B1). Plants with such a trade-off tend to evolve towards the extreme cases, that is strategies where either attractiveness or independent growth is absent (either *α* =0 and *r* _*P*_=1 or *α* =1 and *r* _*P*_=0). A balanced strategy with intermediate investment into both attractiveness and independent growth can only be expected for relatively large values of *s*, which translate into a concave trade-off shape (see supplementary material B). Weinbach et al. (2022) showed that in these cases, the plant could indeed maintain both the mutualistic interaction and selfing or vegetative reproduction and that the pollinator could coexist with the plant for a large range of net growth rate coefficient values (*r*_*A*_). In the following analysis, we therefore chose *s=3*.

Which trait values turn out to be optimal for the plant and its pollinator in a given situation obviously depends on plant and pollinator abundances, as well as on the strength of the mutualistic interaction. For example, with limited pollination service due to low pollinator densities, the plants might invest less into attractiveness (low value of *α*) and more into intrinsic growth (high value of *r*_*P*_). Similarly, with low food availability due to low plant densities and/or low attractiveness, the pollinator might also invest less into the interaction (low value of *β*) and more into other energy sources (high value of *r*_*A*_).

We expect that plant evolution will lead to a value of *α* that maximises the sum of the two positive growth terms in equation 1a, that is *P* ·(*r*_*P*_+ *α β γ* _*A*_ *A*). Analogously, we also expect that pollinator evolution will lead to a value of *β* that maximizes the sum of the two positive growth terms in equation 1b, that is *A* ·(*r ‘* _*A*_ +*α β γ* _*P*_ *P*). For fixed values of *γ* _*P*_ and *γ* _*A*_ the evolution of *α* is then driven by the product *βA* and the evolution of *β* is driven by the product *αP*. Note that the environmental decay as such cannot have a direct effect on the optimal beta, because it does not affect the two growth terms. It does, however, affect the pollinator density *A*, and hence *α*, which in turn also affects the plant density *P* and hence indirectly also *β*. In the face of declining pollinator abundances, both populations will consequently adapt by shifting their energy allocation, which will then either accelerate or decelerate the initial pollinator decline.

### 2.4 Numerical simulation of eco-evolutionary dynamics

Our analysis of adaptive changes in energy allocation of plants and pollinators in response to pollinator decline is inspired by the mathematical framework of Adaptive Dynamics (see Brännström et al. (2013) for a review). Weinbach et al. (2022) performed an in-depth analysis of possible singular strategies in the simplified case without pollinator evolution, including an analysis of how the trade-off shape affects the number and stability of these singularities. The authors derived an analytical expression for the singular strategy for the special case of a linear trade-off, see equation A7 in the corresponding online supplementary material. The result can be transferred to pollinator evolution under the condition that plant evolution is switched off, due to the existing symmetry between plant and pollinator equations in our model.

However, with two evolving traits (here *α* and *β*), analytical calculations using Adaptive Dynamics are typically very complicated (see Dieckmann and Law (1996) for the general method and Kisdi (2006) for an application). The concave energy allocation trade-off adds another level of complexity, meaning that analytical investigations of our model are probably not tractable and hence beyond the scope of this study. Instead, we investigate the eco-evolutionary dynamics of our model using numerical simulations, programmed in Python 3.10. The simulation code is available as online supplementary material.

Each simulation started with a single ancestor plant population and a single ancestor pollinator population at an eco-evolutionary equilibrium. We then performed recurrent “mutation events”. During such a mutation event, we chose one of the populations as “resident” and simply added a new “mutant” population. The traits of the mutant population were chosen from a normal distribution around the parent’s traits. This normal distribution had a standard deviation of 1/2.5758, so that 99% of all randomly chosen numbers were in the interval [-1,1], and was multiplied with Δα and Δβ, respectively, so that Δα and Δβ effectively describe the maximum mutation distance between resident and mutant. The initial per-capita growth rate of the mutant was derived via the so-called “invasion fitness”, that is the change in mutant density over time under the assumption that the mutant *m* is rare and that the resident *r* is at equilibrium densities *P*^*^_*r*_ or *A*^*^ _*r*_, respectively:

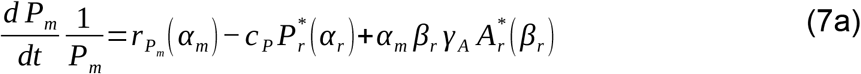

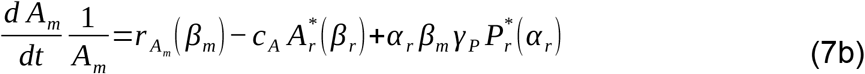

The fate of the mutant population was determined by its traits, which affect the strength of the mutualistic interaction, as well as the state of the environmental decay. The mutant could (i) have a positive growth rate and eventually replace the resident population in case it was better adapted to the current situation, (ii) have a negative growth rate so it quickly went extinct or (iii) coexist with the resident. The latter outcome is known as evolutionary branching and can, in principle, lead to the emergence of complex interaction networks (Brännström et al., 2011; Geritz et al., 1998; Nicolas Loeuille & Loreau, 2005). However, in our system we only observed outcome (i) and (ii).

We used two different implementations of the algorithm outlined above. Following the core hypothesis of the Adaptive Dynamics framework (Brännström et al., 2013; Geritz et al., 1998), the first approach was based on a separation of ecological and evolutionary time scales. In particular, we assumed that mutation events are so rare that the population dynamics of the resident will always reach an ecological equilibrium before the next mutation event takes place. To shorten the simulation runtime, we therefore omitted the calculation of the population dynamics. Instead, we instantly replaced the resident population with the mutant population if the initial per-capita growth rate of the mutant was positive (in line with outcome (i)). By contrast, mutants with negative growth rates were assumed to be not viable and simply removed from the system (in line with outcome (ii)). The code for this first model version thus includes equations (2a) and (2b) for the equilibrium densities, equation (5) for the environmental decay, equations (6a) and (6b) for the trade-off functions, as well as equations (7a) and (7b) for the per-capita growth rates of new mutant populations, as summarised in Fig 1.

**Fig. 1:**
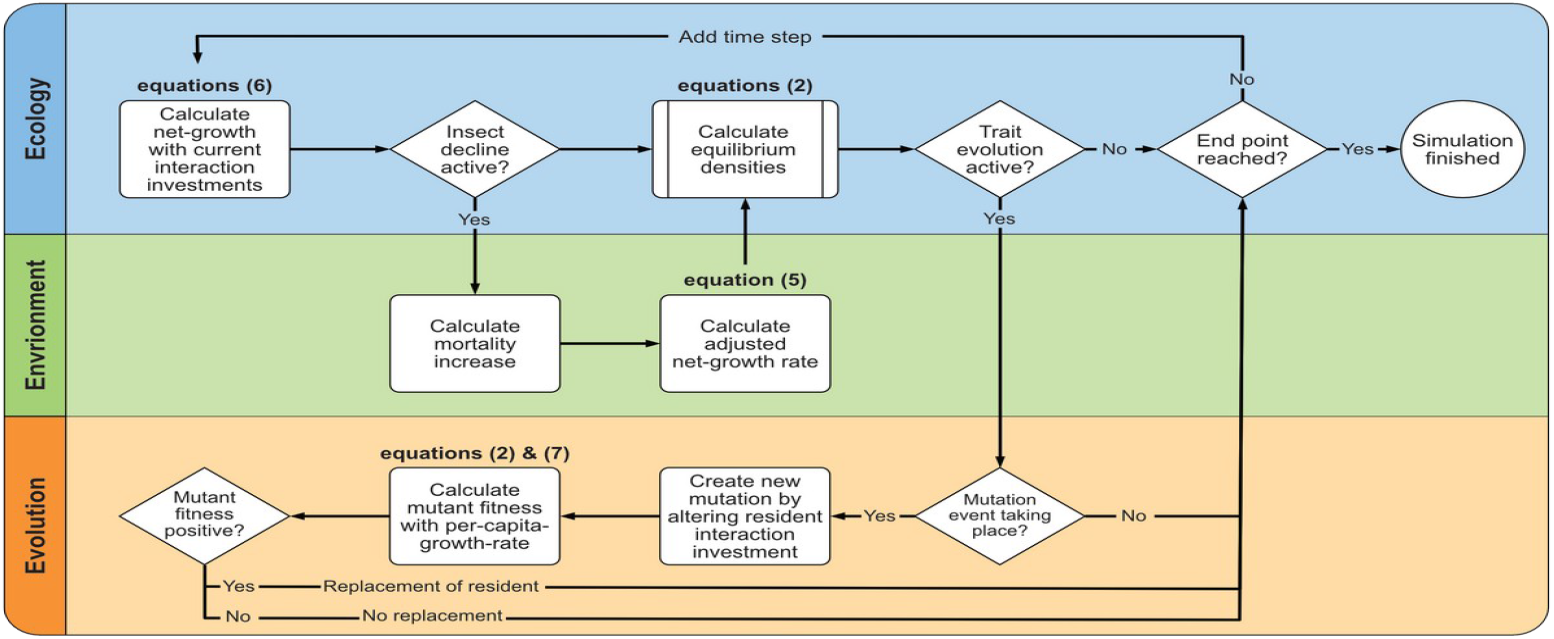
Overview of the model algorithm. The ecological part (in blue) calculates the ecological equilibrium densities, the environmental part (in green) implements the population decline via environmental decay and the evolutionary part (in orange) implements evolution for both the plant and the pollinator populations. Most simulations started at an eco-evolutionary equilibrium and were run with the environmental degradation being active. The end condition was then simply whether or not the pollinator was still alive. Note that the algorithm can also be run without environmental decline, with the goal to pinpoint the initial eco-evolutionary equilibrium. In these cases, the end condition was a fixed time limit. We observed that *α* and *β* typically approached certain values and that new morphs were no longer able to invade as soon as the system was sufficiently close to these values. Evolutionary endpoints that are both convergent stable and evolutionarily stable, as in this case, are known as Continuously Stable Strategies (CSS, see Brännström et al. (2013)). Other types of evolutionarily singular strategies did not occur in our model system.

In a second approach, we then relaxed the assumption of separate ecological and evolutionary timescales. The corresponding algorithm is summarised in supplementary Fig. C1. Instead of instantly replacing the resident with the mutant population, as explained above, we simulated the explicit resident and mutant population dynamics over time. The initial population density of the mutant population equalled the extinction threshold *ε* and was substracted from the resident population. Mutation events were still performed with fixed probabilities *q*_*A*_ and *q*_*P*_, as in the first approach, but only at every 100th time step instead of every time step. The establishment of the previous mutant was typically not completed when the next mutant was added. In consequence, we observed multiple mutant populations of plants and/or pollinators apparently coexisting over time. For computational reasons, at any given time we limited the number of plant and/or pollinator mutants to a maximum of 25 coexisting populations. Population dynamics were implemented using the following set of equations, with indices running over the multiple mutant populations, to accommodate a growing (or shrinking) number of populations:

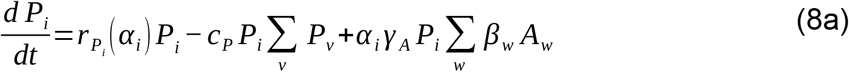

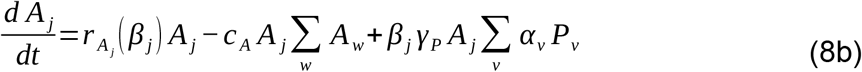

Simulations performed with the first approach were very fast at the cost of limited biological realism. Simulations based on the second approach, on the other hand, were more realistic, but also more time consuming, due to the high number of coupled equations. We therefore performed most of our analysis using onlythe first approach and then repeated part of the analysis using the second one, as a robustness check. In both cases, we compared the extinction time points of pollinators with and without evolution for various parameter combinations, to analyse the impact of eco-evolutionary feedback on pollinator persistence. Additional robustness checks addressed the potential impact of the trade-off shape shape in equation 6 (see supplementary material D), as well as of an alternative functional response that accounts for limiting returns on mutual benefit as partner abundance increases (see supplementary material E).

## 3. Results

### 3.1 Delayed pollinator extinction can occur for high interaction benefits, low competition and slow plant evolution

The pollinator population in our simulations is always doomed to go extinct, due to ever increasing mortality rates, see eq. (5), but the exact time point of extinction depends on the involved eco-evolutionary forces. We found that eco-evolutionary feedback could in principle delay pollinator extinction and thus help those pollinators to survive harsher conditions. This was in particular true for relatively high interaction benefits *γ*_*A*_ *and γ*_*P*_ and low competition *c*_*P*_ and *c*_*A*_ (blue area in Fig. 2). Most of the parameter combinations, however, resulted in self-reinforcing eco-evolutionary feedback that accelerated pollinator extinction (red area in Fig. 2). Note that extremely low interaction benefits and very intense competition led to a rather weak interaction between relatively small populations, which explains why in these cases the effect of eco-evolutionary feedback was less pronounced. Extremely high interaction benefits in combination with very low competition, on the other hand, led to a strong coupling of large populations, which pushed the systems towards unrealistic unbounded growth, as explained in the methods section (see eq. 3).

**Fig. 2:**
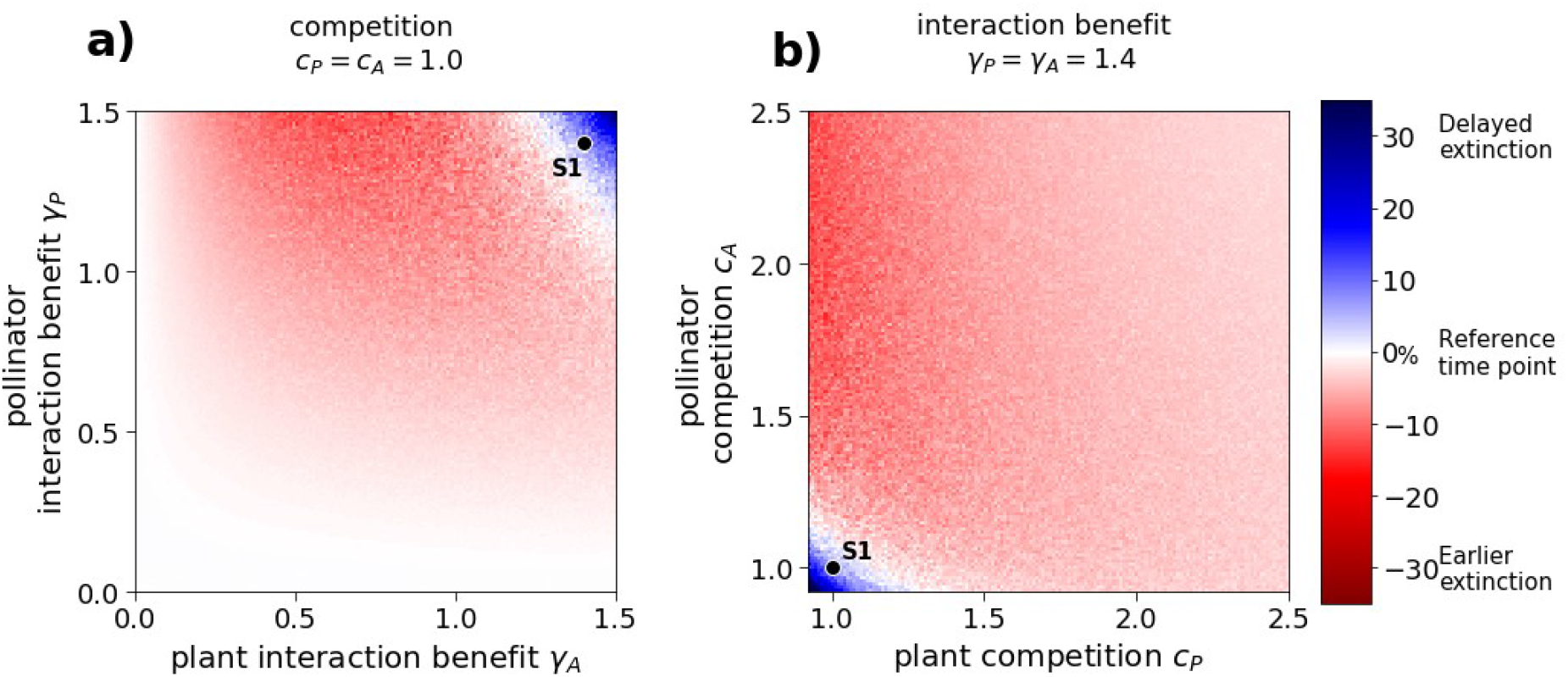
High interaction benefits and low competition can lead to delayed pollinator extinction. The colour code indicates the observed extinction time point for the given parameter combination relative to the extinction time point of the corresponding simulation without evolution in [%]: earlier extinction is shown in red, delayed extinction in blue. White areas indicate no effect of eco-evolutionary feedback on pollinator extinction. Mutation probabilities were *q*_*P*_=10 % for the plants and *q* _*A*_=50 % for the pollinators. All other parameters were chosen according to Table 1, if not indicated otherwise. The time series for the simulation highlighted as S1 can be found in Fig. 5.

Not only the ecological parameters had an effect on the eco-evolutionary forces but also the relative speed of plant versus pollinator adaptation (Fig 3). We typically observed earlier pollinator extinction in the extreme case of only plant evolution (*q*_*A*_*=0, q*_*P*_*>0*, see x-axis in Fig. 3a-c). Delayed extinction was possible in the other extreme case, when plant evolution was switched off, in particular when the interaction benefit was sufficiently high (*q*_*P*_*=0, q*_*A*_*>0*, see y-axis in Fig. 3a-c). In the case of co-evolving interaction investments, we found that delayed extinction became more likely when plant evolution was decelerated or when pollinator evolution was accelerated.

**Fig. 3:**
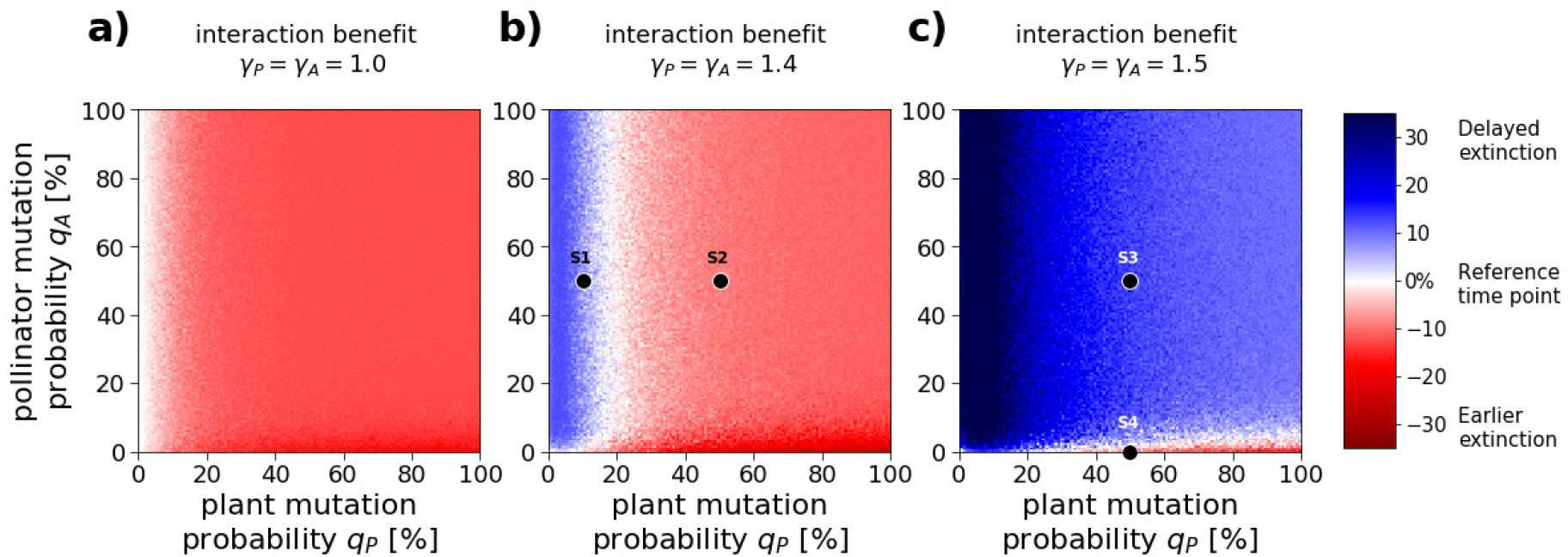
Delayed pollinator extinction is more likely with slow plant and fast pollinator evolution. The colour code indicates the observed extinction time point for the given parameter combination relative to the extinction time point of the corresponding simulation without evolution in [%], as in Fig 2. Parameter values were chosen according to Table 1, if not indicated otherwise. The time series for the simulations highlighted as S1-S4 can be found in Fig. 5.

The effect of evolutionary adaptation on pollinator survival strongly depends on how fast the evolutionary response is in comparison to the environmental decay. In general, we found that delayed pollinator extinction is more likely and/or more pronounced for faster environmental decay, especially if plant evolution is sufficiently slow (Fig 4). In particular, we found that delayed pollinator extinction is possible in a rapidly changing environment, even if pollinator evolution was switched off in the model structure (Fig. 4c).

**Fig. 4:**
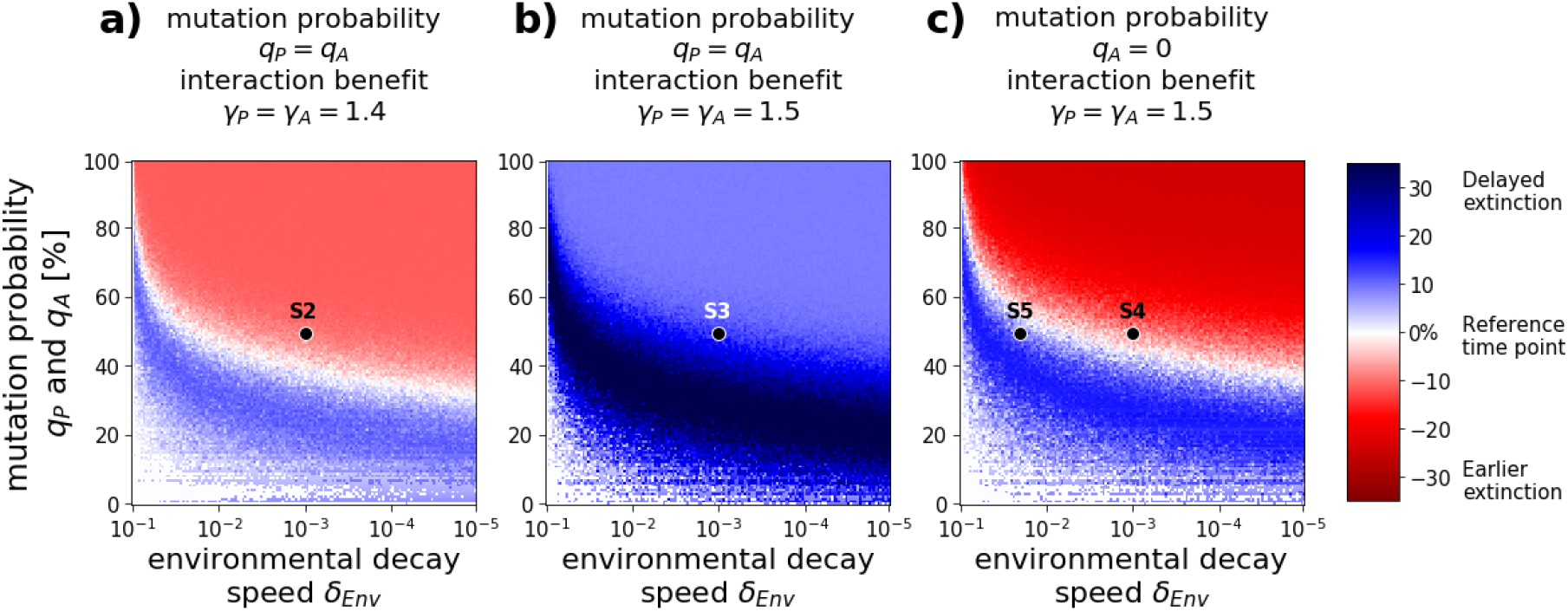
Delayed pollinator extinction is more likely with fast environmental decay. The colour code indicates the observed extinction time point for the given parameter combination relative to the extinction time point of the corresponding simulation without evolution in [%], as in Fig 2. Parameter values were chosen according to Table 1, if not indicated otherwise. The time series for the simulations highlighted as S2-S5 can be found in Fig. 5.

### 3.2 Plants disinvest first, pollinators follow

We always observed the same two processes in our simulations, namely disinvestment of the plants and disinvestment of the pollinators (Fig. 5). Increasing pollinator mortality rates primarily affected the pollinators themselves and resulted in declining pollinator abundances (top row, yellow lines). The plant population then responded to such declining abundances by reducing its investment into the interaction (decreasing values of *α*, third row, green lines) in favour of intrinsic growth (increasing values of *r*_*P*_, not shown). The plant disinvestment slowed down the plant decline that was triggered by the reduced interaction (top row, green line) at the cost of further threatening the pollinator population. The pollinator consequently responded to this double pressure of increasing mortality rates and decreasing interaction strength by also reducing its investment into the interaction (third row, yellow lines). This disinvestment then allowed the pollinator to partly compensate for increasing mortality rates (second row), which in turn also slowed down pollinator decline (top row, yellow lines). The disinvestment of both interaction partners was typically rather slow in the beginning but accelerated during the simulation (third row), which was partly due to the fact that both processes reinforce each other but also reflects the concave shape of the chosen trade-off functions (see eq. 6).

**Fig. 5:**
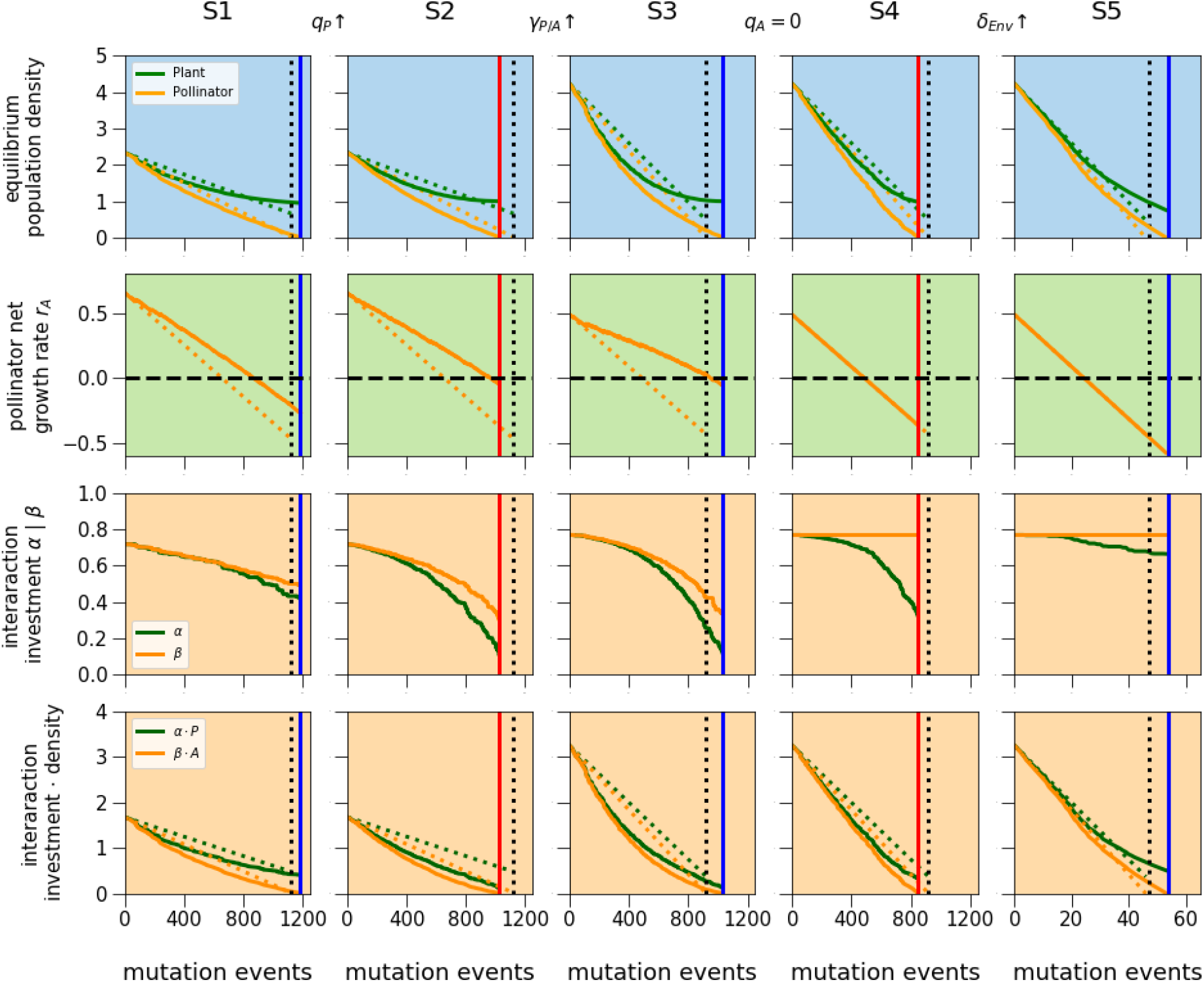
Overview of selected time series. Background colours highlight ecological (blue), environmental (green), and evolutionary (orange) processes. Vertical lines indicate pollinator extinction time points. Note that plant densities always approach 1, which equals the plant carrying capacity *1/c*_*P*_ with *c*_*P*_=1, that is the equilibrium density of the plant in the absence of pollinators. Dotted lines show reference simulation without evolution (*q*_*P*_=*q*_*A*_ =0). Simulation S1 was run with interaction benefits *γ*_*P*_ = *γ*_*A*_ = 1.4 and mutation probabilities *q*_*P*_=10 % for the plant and *q* _*A*_ =50 % for the pollinators. All other parameters were chosen according to Table 1. We then changed one parameter at a time, while keeping all other parameters constant. From S1 to S2: increase plant mutation probability *q*_*P*_=10 % instead of *q*_*P*_=10 %). From S2 to S3: increase interaction benefit (*γ*_*P*_=*γ*_*A*_=1.5 instead of 1.4). From S3 to S4: switch off pollinator evolution (*q* _*A*_=0 % instead of *q*_*P*_=50 %). From S4 to S5: increase speed of environmental decay (*Δ*_*Env*_=0.02 instead of 0.001).

The first process (plant disinvestment) was in most cases harmful for the pollinator, as it weakened the mutualistic interaction, while the second process (pollinator disinvestment) typically helped them to survive longer. Whether or not the combined effect of both processes translated into earlier or delayed pollinator extinction thus strongly depended on their relative strength, which was modified via differences in the mutation frequency (see Fig. 3). In particular, we found that delayed extinction can turn into earlier extinction if plants evolve faster (compare S1 and S2 in Fig 5). However, also the initial situation at simulation start played a key role, which became apparent when analysing simulations with high interaction benefits and/or low competition (see Fig. 2 and 3). These simulations started with a very strong interaction between large populations, indicated via relatively high initial population densities *P* and *A* (see top panel for S3 in Fig 5) and high initial interaction investments *α* and *β* (third panel for S3). Plants then needed more time to actually disinvest, simply because they started at a higher level of interaction investment *α*, so that the mutualistic interaction was maintained longer. At the same time, the beneficial effect of pollinator disinvestment was particularly strong, due to the concave shape of the trade-off function (eq. 6): small mutation steps led to a substantial increase in intrinsic growth *r*_*A*_ as long as the interaction investment *β* was still high, which in turn directly compensated for increasing mortality rates. In consequence, we found that the beneficial effect of pollinator disinvestment typically dominated when the initial interaction was very strong, resulting in delayed extinction, even if both interaction partners had the same mutation frequency (*q*_*A*_*=q*_*P*_ in S3).

The delayed extinction observed in S3 turned into earlier extinction after switching off pollinator evolution, indicating that plant evolution is indeed harmful for the pollinator (S4). However, plant evolution could also be beneficial, in particular when the environmental decay was relatively fast (S5). This surprising result is again due to the shape of the concave trade-off function (eq. 5): small mutation steps led to a substantial increase in intrinsic growth *r*_*P*_ as long as the investment *α* was still high. This had a direct positive effect on plant abundance, which could then indirectly benefit the pollinators as long as the mutualistic interaction was still strong. Note, however, that the positive effect of plant evolution was negligible compared to the negative effect of increased environmental decay, meaning that the pollinators still went extinct much earlier compared to scenarios with moderate or slow environmental decay.

After all these investigations, it remains challenging to predict whether a given parameter combination results in earlier or delayed pollinator extinction, due to the high number of interacting forces that affect trait evolution in our model system. Our simulations suggest that the products *βA* and *αP* can help to synthesise these results. Both products decline over time, independent of whether evolution is taken into account or not (bottom line in Fig. 5). This decline is initially faster when evolution is switched on compared to simulations where evolution is switched off. However, while the speed of the decline remains constant over time when evolution is switched off, it slowly decelerates when evolution is taken into account, meaning that the corresponding curves bend upwards. In some cases, the curves bend fast enough to intersect with the non-evolution-curves before the pollinator density reaches the extinction threshold, meaning that the products *βA* and *αP* are then higher compared to the case without evolution. These are the scenarios that lead to delayed pollinator extinction.

### 3.3 Robustness checks

We acknowledge that the model outcomes presented above are to some extent dependent on the specific trade-off function. In particular, the observed eco-evolutionary dynamics might accelerate or slow down when choosing a higher or lower value of the parameter *s*, which captures the trade-off shape. However, we still find the same qualitative pattern of joint disinvestment if *s* is sufficiently large, meaning that the trade-off function is sufficiently concave. For more detailed information of this matter please refer to the methods section and to the online supplementary Fig. D1 and D2.

Our results are furthermore based on two simplifying assumptions. In particular, we assume that (a) the ecological and evolutionary time scales are separate (see Methods section) and that (b) the mutualistic interaction can be described via a linear functional response (see equation 1). As a final step, we relaxed these assumptions by using a more realistic (but slower) simulation algorithm that captures overlapping timescales and by using a type 2 functional response that takes limiting returns on mutual benefit into account. The results obtained with these model versions were generally consistent with the results presented above but differed in the exact parameter combinations for which delayed extinction could be observed (compare Fig 3 with supplementary Fig. E1 and E2).

## 4. Discussion

In this study, we investigated how eco-evolutionary dynamics within a mutualistic two-player system shape the system’s response to one-sided population decline. Our model analysis revealed two important key insights. First, in all scenarios that were studied a joint disinvestment of both interaction partners in response to the decline of one interaction partner was found. Second, despite the generality of the first key finding, it still remains difficult to predict whether such joint disinvestment eventually translates into delayed or earlier pollinator extinction. This is due to a highly non-trivial interplay between the ecological, environmental and evolutionary processes considered in the model, as well as the relative speed of these processes. In summary, we found that delayed pollinator extinction was possible in cases where the products *βA* and *αP* remained relatively high, that is in cases with sufficiently high interaction benefits, sufficiently low intraspecific competition and relatively slow plant evolution.

The observed joint disinvestment is surprising at first, because one might naively assume that at least one interaction partner would invest more into the interaction, not less, in order to maintain its benefits. However, the disinvestment can be easily explained as a direct consequence of the assumed energy allocation trade-off. This trade-off introduces a cost to the mutualistic interaction, meaning that maintaining interaction benefits goes hand in hand with losing other energy sources, such as alternative food sources, reproduction spots or shelter. These alternative resources become relatively more important as soon as the mutualistic interaction is weakened due to pollinator decline, leading to a shift in energy allocation in favour of those other resources, which in turn further weakens the interaction.

Those scenarios in which joint disinvestment led to delayed pollinator extinction provide potential for evolutionary rescue (see Gonzalez et al. (2013) or Bell (2013, 2017)): If the pollinator mortality rate does not continue to increase forever, as in our study, but instead reaches at a value that is just high enough to push the pollinators below the extinction threshold in the absence of evolution, then those pollinators might still be able to survive when evolution is switched on, due to eco-evolutionary feedback. Note that in the past, evolutionary rescue theory has mostly been applied to single populations (see for example the summarising reviews of C.

Carlson et al. (2022) and Bell (2017)), while our model demonstrates that the concept of evolutionary rescue can be extended towards multi-species systems: The survival of a target species does not only depend on its own evolutionary trajectory, but also on adaptive responses of their interaction partners. This is in line with findings by Yamamichi and Miner (2015), who showed that a non-evolving predator population can be rescued from extinction by adaptive evolution of its prey, resulting in indirect evolutionary rescue. It is also in line with recent work by (Yacine et al., 2021), who analysed body mass evolution within model food webs. Their simulations suggest that long-term biodiversity losses triggered by warming are considerably higher when evolution is slowed down or switched off completely, pointing towards evolutionary rescue at the community scale.

However, most scenarios that were studied actually resulted in accelerated pollinator extinction, which provides potential for evolutionary murder instead of evolutionary rescue (as suggested by Parvinen (2005)): If pollinator mortality rates increase only until they reach a value that is still smaller than the critical value at which pollinators go extinct in the absence of evolution, then pollinator survival would nevertheless be threatened when evolution is switched on, again due to eco-evolutionary feedback. These scenarios consequently challenge the naive assumption that evolution is generally beneficial for species survival and coexistence, in the sense that it will help ecosystems to adapt to external stressors. This insight is consistent with previous studies on evolutionary trapping and suicide (see for example Ferriere and Legendre (2013) or Uchiumi et al. (2023)), evolutionary extinction debts (Norberg et al., 2012) and even eco-evolutionary tipping points (Dakos et al., 2019), including the original model version without pollinator evolution (Weinbach et al., 2022).

We acknowledge that due to its simplicity, our model clearly does not capture the full complexity of plant-pollinator interactions. This is in particular true for the linear functional response that we chose to represent the mutualistic interaction. However, the goal of this study was not to model a specific mutualistic system with all its typical characteristics. Instead, we focus on the impact of the eco-evolutionary feedback loop on the evolution of the investment into mutualism and on pollinator persistence. We argue that whether or not this feedback loop is self-reinforcing or self-dampening depends on whether or not the biotic interaction is mutualistic or antagonistic, as well as on the strength of this interaction, but not on the exact shape of the underlying functional response. This is not only corroborated by our robustness check (see supplementary Figure E2) but also in line with a recent review of models for pairwise mutualistic interactions, which showed that population dynamics of mutualisms are qualitatively robust across derivations, including levels of detail, types of benefit, and inspiring systems (Hale & Valdovinos, 2021).

Because our model is quite abstract and general, it could, at least in principle, describe any type of mutualistic interaction experiencing one-sided population decline. However, we deliberately chose to study our model through the lens of plant-pollinator systems, given that our results are of direct relevance for understanding and predicting the consequences of global pollinator decline (Po tts et al., 2010). Our results offer two contrasting hypotheses for eco-evolutionary responses of plant-pollinator interaction networks to insect decline. Both hypotheses are based on the assumption that our model system represents a small network module embedded into a larger network but differ in the interpretation of plant and pollinator growth rates. If these growth rates are interpreted as energy gain from interaction with alternative partners, that is with the extended network in which the plant-pollinator module is embedded, then interaction disinvestment directly translates into evolution towards generalism or towards specialisation on alternative interaction partners. In consequence, we might expect to see network rewiring, but not necessarily network collapse. On the other hand, disinvestment can also be interpreted as evolution towards self-reproduction, in particular on the plant side, with potentially detrimental long-term consequences for the whole system due to reduced gene flow and inbreeding depression (Lepers et al., 2014).

The goal of this study was to see if the reinforcing ecological feedback loop observed between two mutualistic species still occurs when coevolution of both mutualistic partners is considered. We saw here that evolution, via the joint disinvestment of both partners, speeds up the decline of the already threatened population. Species will evolve away from interacting with bad quality interactors in the presence of energetically more efficient alternatives (Werner et al., 2018), given that mutualism is always a service at a cost of other reproduction and survival means (Bronstein, 2001). Our model is useful in identifying interactions that might be too costly to maintain in future scenarios. Specialist species are already known to be more prone to extinction (Biesmeijer et al., 2006), but according to our model this is also the case for species who provide a bad mutualistic service (e.g. not efficient pollinators) and suffer already from strong internal competition (e.g. competition for nesting sites in very artificial areas or for partners in very fragmented ones). Thus, if restoration attempts are made to rescue endangered mutualistic populations, the most fragile ones would be those populations with inefficient pollinators and/or high intrinsic competition.

However, because evolution reinforces any tendency, it could also accelerate any recovery that could be implemented with restoration attempts, which is motivating for conservation actions. According to our model, restoration should focus not only on the endangered species (e.g. by providing more food, nests and potential reproduction partners) but also on the species interactors. Improving the plant’s living conditions could also decrease its internal competition and improve its interaction gains, reducing the speed of disinvestment into the interaction or even reversing the evolutionary disinvestment once the energetic balance is again in favour of mutualism.

## Acknowledments

We thank all members of the Eco-Evolutionary Modelling group at the University of Hohenheim for many inspiring discussions throughout the whole project as well as valuable feedback on the final manuscript. We furthermore thank Fynn Bachmann for assistance with programming.

## Ethics approval

Not applicable.

## Consent to participate

Not applicable.

## Consent for publication

Not applicable.

## Availability of data and material

Not applicable.

## Code availability

The simulation code will be made available as supplementary material.

## Author Contributions

KTA and FW designed the study, with minor contributions from AW and CZ. FW carried out the numerical simulations and analysed the data. KTA took the lead in supervision and project organisation. All authors discussed the simulation results. FW and KTA wrote the first manuscript draft, with input from AW. All authors contributed critically to revisions. All authors read and approved the final manuscript.

