## Supplementary material for "Eco-evolutionary dynamics in two-species mutualistic systems: One-sided population decline triggers joint interaction disinvestment"

### - online supplementary material -

**Franz Weyerer**[1]
**Avril Weinbach**[2]
**Christiane Zarfl**[1]
**Korinna T. Allhoff**[1,2*]

1) Eberhard Karls Universität Tübingen, Tübingen, Germany
2) Universität Hohenheim, Stuttgart, Germany

**Corresponding author:** Dr. Korinna Theresa Allhoff, University of Hohenheim, Institute of Biology, FG Eco-Evolutionary Modelling (190m), Garbenstr. 30, 70599 Stuttgart, Germany.

**ORCIDs:** Allhoff (0000-0003-0164-7618); Zarfl (0000-0002-2044-1335); Weinbach (0000-0001-7598-1112)

**Content:**

A. Mathematical derivation of the coexistence fixed point

B. Additional information about the trade-off shape

C. Overview of the evolutionary algorithm with population dynamics

D. Robustness check 1: More or less concave energy allocation trade-off

E. Robustness check 2: Overlapping ecological and evolutionary time scales with or without saturating functional responses

### A) Mathematical derivation of coexistence fixed point

Here we present the analytical derivation of the equilibrium population densities presented in equation 2 in the main manuscript. The coexistence fixed point of our dynamical system (see equation 1 in the main manuscript) can be found at the intersection of the non-trivial isoclines, which can be calculated by setting the per-capita growth rates to zero:

$$\frac{dP}{dt}\frac{1}{P}=r_P-c_P P+\alpha\beta\gamma_A A=0$$

$$\frac{dA}{dt}\frac{1}{A}=r_A-c_A A+\alpha\beta\gamma_P P=0$$

Solving these equations for *P* and *A* results in:

$$P=\frac{r_P+\alpha\beta\gamma_A A}{c_P} \quad \text{(A1)}$$

$$A=\frac{r_A+\alpha\beta\gamma_P P}{c_A} \quad \text{(A2)}$$

Both equations have to be fulfilled for the system to be at equilibrium. The equilibrium density *A** of the pollinators can consequently be found by inserting the equation for the first isocline (A1) into the equation of the second (A2):

$$A^*=\frac{r_A+\alpha\beta\gamma_P\frac{r_P+\alpha\beta\gamma_A A^*}{c_P}}{c_A}$$

$$A^*=\frac{r_A+\alpha\beta\gamma_P\frac{r_P+\alpha\beta\gamma_A A^*}{c_P}}{c_A} \qquad \cdot\frac{c_P}{c_P}$$

$$A^*=\frac{r_A c_P+\alpha\beta\gamma_P r_P}{c_A c_P}+\frac{\alpha^2\beta^2\gamma_A\gamma_P A^*}{c_A c_P} \qquad -\frac{\alpha^2\beta^2\gamma_A\gamma_P A^*}{c_A c_P}$$

$$A^*\left(1-\frac{\alpha^2\beta^2\gamma_A\gamma_P}{c_A c_P}\right)=\frac{r_A c_P+\alpha\beta\gamma_P r_P}{c_A c_P} \qquad \div\left(1-\frac{\alpha^2\beta^2\gamma_A\gamma_P}{c_A c_P}\right)$$

$$A^*=\frac{r_A c_P+\alpha\beta\gamma_P r_P}{c_A c_P}\frac{1}{1-\frac{\alpha^2\beta^2\gamma_A\gamma_P}{c_A c_P}}$$

$$A^*=\frac{r_A c_P+\alpha\beta\gamma_P r_P}{c_A c_P-\alpha^2\beta^2\gamma_A\gamma_P}$$

Analogously, the equilibrium density *P** of the plants be found by inserting the equation for the second isocline (A2) into the equation of the first (A1):

$$P^*=\frac{r_P+\alpha\beta\gamma_A\frac{r_A+\alpha\beta\gamma_P P^*}{c_A}}{c_P}$$

$$P^*=\frac{r_P+\alpha\beta\gamma_A\frac{r_A+\alpha\beta\gamma_P P^*}{c_A}}{c_P} \qquad \cdot\frac{c_A}{c_A}$$

$$P^*=\frac{r_P c_A+\alpha\beta\gamma_A r_A}{c_A c_P}+\frac{\alpha^2\beta^2\gamma_A\gamma_P P^*}{c_A c_P} \qquad -\frac{\alpha^2\beta^2\gamma_A\gamma_P P^*}{c_A c_P}$$

$$P^*\left(1-\frac{\alpha^2\beta^2\gamma_A\gamma_P}{c_A}\right)=\frac{r_P c_A+\alpha\beta\gamma_A r_A}{c_A c_P} \qquad \div\left(1-\frac{\alpha^2\beta^2\gamma_A\gamma_P}{c_A c_P}\right)$$

$$P^*=\frac{r_P c_A+\alpha\beta\gamma_A r_A}{c_A c_P}\frac{1}{1-\frac{\alpha^2\beta^2\gamma_A\gamma_P}{c_A c_P}}$$

$$P^*=\frac{r_P c_A+\alpha\beta\gamma_A r_A}{c_A c_P-\alpha^2\beta^2\gamma_A\gamma_P}$$

### B) Additional information about the trade-off shape

We assume the following functions to describe the trade-off in energy allocation for plants and pollinators:

$$\left(\frac{r_P}{r_{max}}\right)^{s_P}+\left(\frac{\alpha}{\alpha_{max}}\right)^{s_P}=1 \qquad \text{and} \qquad \left(\frac{r'_A}{r_{max}}\right)^{s_P}+\left(\frac{\beta}{\beta_{max}}\right)^{s_P}=1$$

These equations can be solved for the respective growth rate*:*

$$\left(\frac{r_P}{r_{max}}\right)=\left(1-\left(\frac{\alpha}{\alpha_{max}}\right)^{s_P}\right)^{1/s_P} \qquad \text{and} \qquad \left(\frac{r'_A}{r_{max}}\right)=\left(1-\left(\frac{\beta}{\beta_{max}}\right)^{s_P}\right)^{1/s_P}$$

With the assumption of $r_{max}=1$ and $s_P=s_A=s$ these equations finally simplify into the trade-off functions presented in the main manuscript (see equation 6):

$$r_P=\left(1-\left(\frac{\alpha}{\alpha_{max}}\right)^{s}\right)^{1/s} \qquad \text{and} \qquad r'_A=\left(1-\left(\frac{\beta}{\beta_{max}}\right)^{s}\right)^{1/s}$$

Note that the exponent *s* captures the shape of the trade-off function, as visualised in Fig. B1.

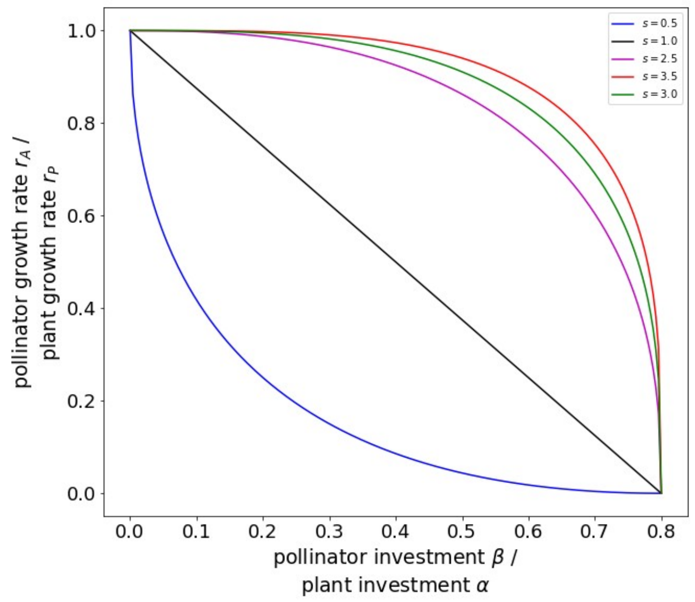


***Figure B1:*** *The shape of the trade-off function depends on the value of the exponent s. With s=1, the trade-off simplifies into a linear function (black line). Values of s<1 result in a convex trade-off (blue line), while values of s>1 result in a concave trade-off (pink, green and orange lines). The simulation data presented in the main manuscript was based on s=3. Robustness checks were performed with s=2.5 and s=3.5, as shown in supplementary Fig. D2.*

'

### C) Overview of the evolutionary algorithm with population dynamics

The algorithm of our simulations is always structured in three parts: "ecology" (blue), "environment" (green) and "evolution" (orange). The ecological part was simplified for the investigations presented in the main text. There, we omitted the explicit calculation of population dynamics by simply replacing the resident with the mutant for the purpose of reducing the simulation runtime. Fig. C1 shows the extended algorithm with explicit calculation of population dynamics that we used for the robustness checks in Fig. E1 and E2.

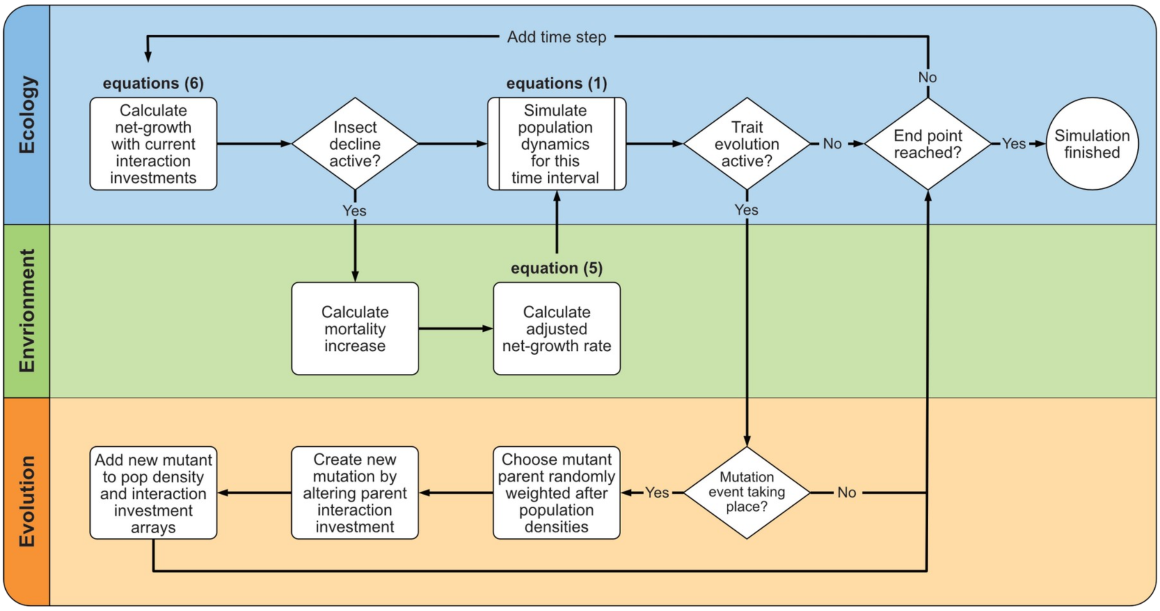


*Figure C1:* ***Overview of the extended model algorithm for overlapping timescales****. The ecological part (in blue) calculates explicit population dynamics, the environmental part (in green) implements the population decline via environmental decay and the evolutionary part (in orange) implements trait evolution for both the plant and the pollinator. Simulations typically started at an eco-evolutionary equilibrium and were run with the environmental degradation being active. The end condition was then simply whether or not the pollinator was still alive. Note that the algorithm can also be run without environmental decline, with the goal to pinpoint the initial eco-evolutionary equilibrium. In these cases, the end condition was a fixed time limit, as in the simplified algorithm described in Fig 1.*

### D) Robustness check 1: More or less concave energy allocation trade-off

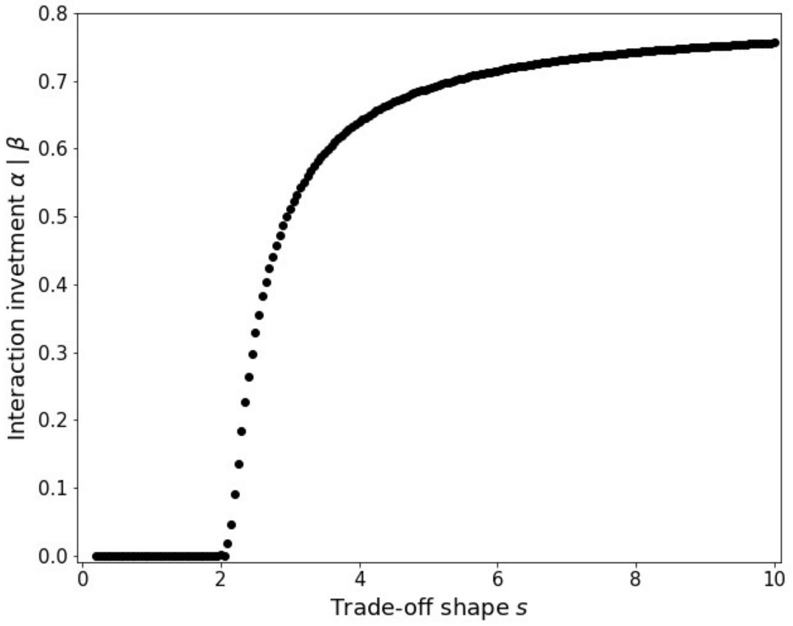


***Figure D1:*** *The parameter s regulates not only the trade-off shape (as shown in supplementary Fig B1) but consequently also the eco-evolutionary endpoint of the dynamics within our model. A mixed energy allocation strategy where plants and pollinators invest both into intrinsic growth and into the interaction with the partner is only possible for values of s>2. In these cases we observed that the system approached an evolutionary endpoint that is both convergent stable and evolutionarily stable (a so-called CSS) as long as the pollinator decline was deactivated. Parameter values:* $c_P=c_A=\gamma_P=\gamma_{A.}=1$ *and* $\delta_{Env}=0$.

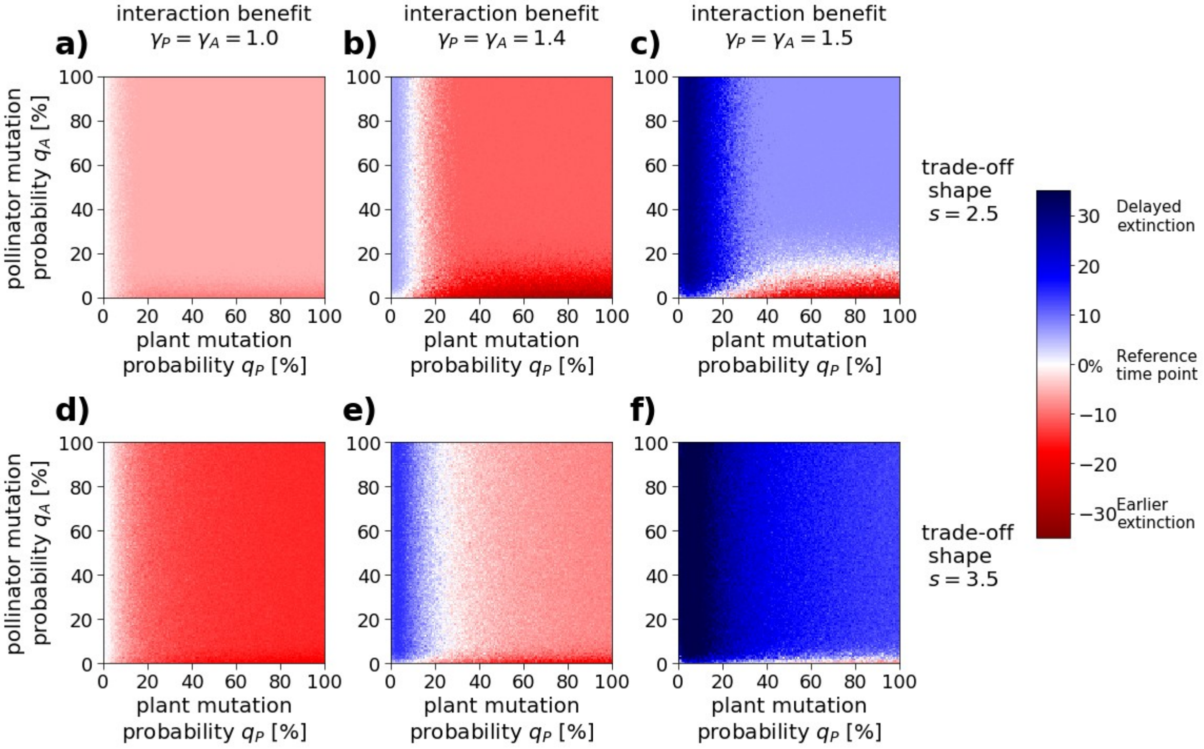


***Figure D2:*** *Same analysis as in Fig 4 but with s=2.5 (less concave trade-off ) and s=3.5 (more concave trade-off) instead of s=3 (see supplementary Fig B1 for comparison). In all three cases, we observed the same qualitative pattern of joint interaction disinvestment. Only the threshold values between areas in the parameter space that led to earlier versus later extinction shifted slightly. Overall, delayed extinction tended to be more likely and/or more pronounced in simulations with a more concave trade-off.*

### E) Robustness check 2: Overlapping ecological and evolutionary time scales with or without saturating functional responses

We relaxed some of our model assumptions with respect to separate time scales and linear functional responses. Overlapping timescales were implemented using the algorithm described in supplementary material C. Type-2 functional responses were implemented by modifying equation 8 into

$$\frac{dP_i}{dt}=r_{P_i}(\alpha_i)P_i-c_P P_i\sum_v P_v+\alpha_i\gamma_A P_i\frac{\sum_w \beta_w A_w}{1+\sum_w \beta_w A_w} \quad (8a)$$

$$\frac{dA_j}{dt}=r_{A_j}(\beta_j)A_j-c_A A_j\sum_w A_w+\beta_j\gamma_P A_j\frac{\sum_v \alpha_v P_v}{1+\sum_v \alpha_v P_v} \quad (8b)$$

We found that the results obtained with overlapping timescales were generally consistent with the results presented in Fig 3 in the main text. They differed, however, in the exact parameter combinations for which delayed extinction could be observed, as summarized in Fig. E1. In particular, we found that the critical value of the interaction benefit $\gamma$ at which the system moved from delayed to earlier extinction tended to be lower with overlapping timescales compared to the simplified model version. We furthermore observed slightly smaller effect sizes, in particular concerning earlier extinction. We explain these differences via the time lag between the 'optimal' evolutionary strategy of the plants, as predicted based on the assumption of separate time scales, and the evolutionary strategy observed during simulations with overlapping timescales. In the latter case, the plants simply did not have enough time to fully adjust their energy allocation strategy to the current environmental condition, meaning that the mutualistic interaction was maintained longer, which in turn lessened the detrimental effect of plant evolution on pollinator survival.

The critical value of the interaction benefit γ was finally even lower when saturating functional responses were taken into account in addition to overlapping time scales. With saturation, the mutualistic interaction was less beneficial, which explain lower initial investment and also reduced initial densities at simulation start. This led to overall shorter simulations since the pollinators were already closer to extinction, meaning that plants had less time to actually disinvest.

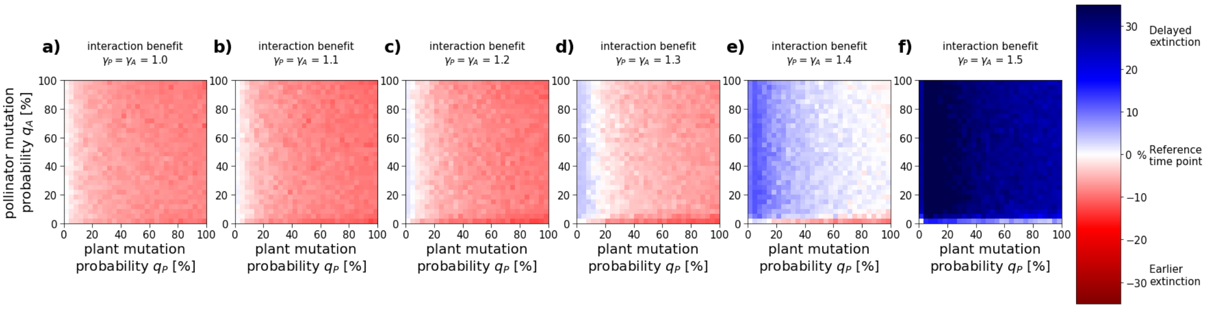


***Fig. E1: Heatmaps with overlapping timescales and intraspecific variability in resource allocation***. *The colour code indicates the observed extinction time point for the given parameter combination relative to the extinction time point of the corresponding simulation without evolution in [%], as in Fig 2. Environmental decay speed was* $\delta_{Env}$=0.00001 *and mutation events took place every 100 time steps. This parameter choice is consistent with a decay speed of* $\delta_{Env}$=0.001 *used in Fig 3, given that Fig 3 is based on a simplified evolutionary algorithm, where mutation events happen at every single time step. Other parameter values were chosen according to Table 1, if not indicated otherwise.*

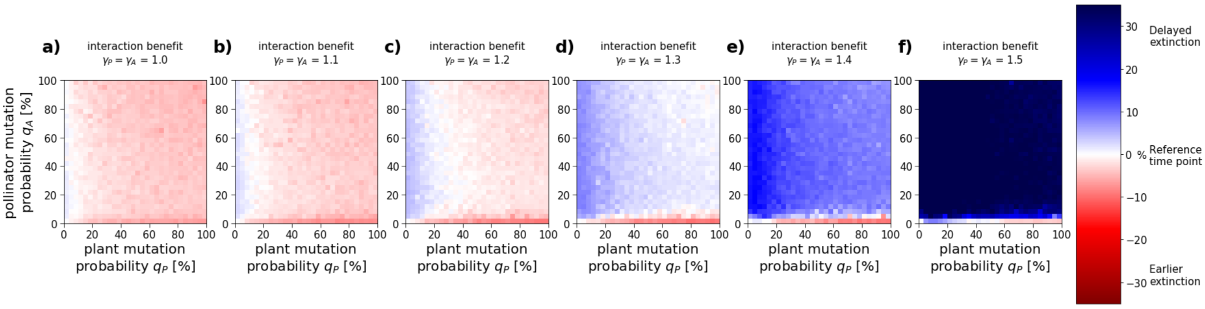


***Fig. E2: Heatmaps with overlapping timescales, intraspecific variability in resource allocation and type-2 functional responses***. *Parameters were chosen as in Fig E1 or according to Table 1. The only difference between Fig E1 and E2 is the saturation of the functional response, which accounts for limiting returns on mutual benefit as partner abundance increases.*
